## Supplemental Sequences for "A protocadherin mediates cell-cell adhesion and integrity of the oral placode in the tunicate Ciona"

**Supplemental Sequences File, Vedurupaka et al.**

**(some sequences are not validated by whole-plasmid sequencing yet)**

Protocadherin.e *in situ* probe template (from Gibboney et al. 2020):

TAAGTGCACTGAGTTGTGTAAAACATATGGACACTGTGATACGTGCTGGATGCCAAGCTCAGGTTATCCCGTGGAGCAGACGAGCCAAATTCCAAATGACGTCACAACACCATGGCAACCGAAGTTCTCCGCCACGAGGAAGGATAGCGGATGTTCTGTCGAAACTATGTCGTCCGCGCTCTCTTATTACAAGCTGACGCGGTCTGAGCAACAGCGACGTCGTAATGATGACACACTTCCAGACCTTGTAGCGAAGGGGCGTACAGCCATGGGCAGTGACGCTAATTCTTTGATGTCTTCAACAAACTCTTCAGGAGCGTCATCGACACGCACCGGTGACCTTAAACATACGTCATCAATTCAAAAATCGCCAACTCATCGGGCTTTGAAAAAGTCCAACAGCGATATAGGGTATGAAGCGAAAGACGCAACACCACGGCAGGTCGTTATTCCTAATAGCCTATCCACACAATGCTAAAGTCGCCTCCCAGGTTTTCCAT

Six1/2 promoters (adapted from Abitua et al. 2015):

>Six1/2 -4285/-1 (used to drive H2B::mCherry)

gtgacaggaaacgtctaggtttaaaacgtatttattgacttctcacttttccggtgatggttttgatttcatttacacgttttggtgcttagctatcttgtagcacggtttagtgtgatacgcataaacgggttgttgaaagtgatgctttgtgctaaacaatgtacgcgggatgtcttagtcacacgcacgtttagatcttgccgctgttagcaccttatgttactaacataatgtttatccataggatcaaagggaacgcttagtgttttacaaataacaaaatatcgtttcactgggaattttatttgtatcctaatggaaaaagtcattttataggaaaaaatctaaccactttatataattatagagggggaacgtgatagcgtttctacaccgagcattgctttgatattgagccacgtcagtaacgaacgatttgaaattaaattaaaacaatttgtttacggttgacaccaagcgcgattgatgttcactctcaggacacttcataatatgtcgcggcctacaacaacattaccatggctgtgaaatgaacagttgacttttaacatctttcagtcgtctggttaaaaaaaggagtcaaccggtacaagattagtttagtttctgacactaatacgtcacactcagttaacgttgacaacgattcaatgacgtatcacgggtcggtcatggtcgagtggaaatatggggaactcaattctgcaaagtggatttctggattatgactcaatccgtaccgacaaggtgttacatgacaggaaaatataaaaatgtaaacagtctgaccgcgttttaaacgtaggattgctctttataacgatgggaattaaagctgctgctggtagaattccccacggtagttttgtttgagaaaggtttgaattggcaattaggcgtgtgaacgtttgagtaaaacatggatagaacgatttattgtgagtcgtgttttggctactgaaatcaaagtatttatcaccgacttaagtgattgatgtgtttgtttgaaatgtggatccccgcaccatttacaataagttataatgatcatgacacggataagtatttaacacttctcaaaatactgtgagttaagtaaaacttcaaagcgttttctgaaaaaaagagttatctataccaactttactattaaaacatcttgtgacaatggcaaacatagggggagtttgttaatacatgcagaccagttgcggttaggttttcgcatttgtgtatggtattaggtttaatgcgtatactattttatatcatgatactatatcattttgtacaatatcatgttgtcgtaaaagtgggaaggattatatagcttgccttgtgtaaaaacatggggtgcggggtggccaagacccaatgcaccatcggctttgatgaatcacgttaattaggggctacgtgggaaagaggggtgggcagagaaaaatctgtgggaaaatgattcgtcttcacgacccagcgcccccgataagacagataaaggtagaacgtcgctcacactctgctacctttgcactgacgagcgataacgagtttttctctcagtggaaaccagcgaaaagttaagaaatagacagtgaccgaaagtgcattaactagtttgctattacattctattatttgtttaacttttgtttttatcgaaacatggacgctgaacttcgattagtgtgtgacgtcactcaacacactgtgatctaaaaattggcaacttcctttttcgtctcgaagtgttcgccctcgtgtctctcttggtctaacacacgcaatagttgaaataagtgagcatctgataaatttgttagaataagaacttagcaatgaagtggaaaaagcgaaaacaatggtttatccgaggtagtagagtagatttctaagcaacacagaaagtttgtaacaacaatgaaagaatattaaacgcactgagattattgtaattgtagtaacatacctttcacattttcaaacgatctgtcaactgctaaactgccaattaatattaacgcatatttgcatagcaattatttactaattccgacacgcaaggaccgatttcccaacgctgtttgatcaaagggaatcgctttgcttgtttaatgcgatcatcaacgctacaaatgtcagcaatcggatattttaagctccgtattaagtcacgtgctacgagcgtgatgaagtggaacgagttgtttacttttaaactcacgtggcgttaattggtcggattaaagtgcgcgtacatgtaagaaaacaattgtctattgtttttcttataattttgtctataaaagcgggaactatcaaaagtatttgttaaccactaaattaaagatatcacgtaacggaaaactggattaataaaacttcaaagattttaaaaagttccgttctataacaacaaaagtgggaaattaacgaataaaagctaagtaatttaaataggaggtataatttttaaaatagtcaaacttgacaagctatagataccaaattagttagaagagaatagcgatacttttaaattgttaggtgcgttgaaatatttgtataacattgattttgccatgggtggtaataaaaccgtttaaaacctttaaggtatgttttcaagcttttatatatattgtaaatgatgtgtttgtttactgaactgcattaaacattcttggattgcattcttatatatttagtatagggtgggggaagatggaagacctttttgatattttctcgtctcatctggtggtaaacaaagaaaattcaattaattataaaaccgtatcctcacaactccaatagatcgttgatatttgtttaaaacacgatcaggctatttggatattatgcactaaaggtgtcccgtcttcccccactctttctactatatacagtggggtgggagaccatggaaaagtttttcattgtattttctcgtccaatctgccagtaaacaagaaaaaacattgaaaaaaaataatatttcgtattctcacgacttccatagaccgtagaatacaatcagaatattttaatattatatactaaaggtgtcccatctccctcacagtactgtatatgagtagtgttttgataagcgataaaatggcgggtatgatgagtattggagcgtcgatcgacaaaatatcgttgcatgcattatacataaggctgaaaccaccacatgacgtttttacaagaggaaaaataacgaaaccctctggtctatgtgcagatctaccaaaatcagcaacgataaagcaatttgttcatgcctatgcatctcctattaatgtagtataagccgaggatacactgcaatcattatacataaatgctctttttgaattataatcacggttaccttgtcaacgcgttcgaaagctttgtttaatattctaacaggtggcgcggtggagcgaatacgtttgttaatcattggtgtatgagcgacactttagggtgttggttatttcaatatgatagcatatatcatacacgataccaacattaaaaatgggtttttagtgtaatttatttgtaaatcgaacgtattcctaaaattataccctaattgttcataattgcaacgtatgcaacgattaaaagttggagttttaagtcaaaaagaaatgaatagcttagaaggtggaagattaaaaataaagtgaaacgtatgtttttgaagagtgttgtctgccaaccttgtgcccttacgtgaaaaaagagattcgaaaacgagagaacttgattgatcggctggtgacatccgacagtagaatctcaaatcagaatctgtttttcgtgtcttctcgcgcactccttattaatgtgttagattcaattttaagcgacagtagtttgggtgaatgtacacctgcgactgatgttactgaaccactttgtttaaataaaggaggaagagaacagaaatagactttaattattatcgcatgtagtcaggttttgcggtcagattgagtctttaatttgtgtttgtattaaattaacactggttgtgtagcaatgtactaaaaaataccctcagatcaaatatattgctgaaatccgtcgccattttagttttatcagttccaccattaacgttggatgataatgttaatttgggaaataaattttatatttttcattttagaactgagtaaaacacttggagtgttctgcacttaacgaggttttcatcgtaacggaaacacattattctacaatcacggcctcagagcca

>Six1/2 -3047/-4 (used to drive CD4::GFP)

ggttttcgcatttgtgtatggtattaggttcaatgcgtatactattttatatcatgatactatatcattttgtacaatatcatgttgtcgtaaaagtgggaaggattatatagcttgcctagtgtaaaaacatggggtgcggggtggccaagacccaatgcaccatcggctttgatgaatcacgttaattaggggctacgtgggaaagaggggtgggcagagaaaaatctgtgggaaaatgattcgtcttcacgacccagcgcccccgataagacagataaaggtagaacgtcgctcacactctgctacctttgcactgacgagcgataacgagtttttctctcagtggaaaccagcgaaaagttaagacatagacagtgaccgaaagtgcattaactagtttgctattacattctattatttgtttaacttttgtttttatcgaaacatggacgctgaacttcgattagtgtgtgacgtcactcaacacactgtgatctaaaaattggcaacttcctttttcgtctcgaagtgttcgccctcgtgtctctcttggtctaacaaacgcaatagttgaaataagtgagcatctgataaatttgttagaataagaacttagcaatgaagtggaaaaagcgaaaacaatggtttatccgaggtagtagagtagatttctaagcaacacagaaagtttgtaacaacaatgaaagaatattaaacgcactgagattattgtaattgtagtaatatacctttcacattttcaaacgatctgtcaactgctaaactgccaattaatattaacgcatatttgcatagtaattatttactaattccgacacgcaaggaccgatttcccaacgctgtttgatcaaagggaatcgctttgcttgtttaatgcgatcatcaacgctacaaatgtcagcaatcggatattttaagcttcgtattaagtcacgtgctacgagcgtgatgaagtggaacgagttgtttacttttaaactcacgtggcgttaattggtcggattaaagtgcgcgtacatgtaagaaaacaattgtctattgtttttcttataattttgtctataaaagcgggaactatcaaaagtatttgttaaccactaaattaaagatatcacataacggaaaactggattaataaaacttcaaagattttaaaaagttccgttctaacaacaaaagtgggaaattaacgaataaaagctaagtaatttaaataggaggtataatttttaaaatagtcaaacttgacaagctatagataccaaattagttagaagagaatagcgatacttttaaattgttaggtgcgttgaaatatttgtataacattaattttgccatgggtggtaataaaaccgtttaaaacctttaaggtatgttttcaagcttttatatatattgtaaatgatgtgtttgtttactgaactgcattaaacattcttggattgcattcttatatatttagtatagggtgggggaagatgtaacacctttttgatattttctcgtctcatctggtggtaaacaaagaaaattcaattaattataaaaccgtatcctcacaactccaatagatcgttgatatttgtttaaaacacgatcaggctatttggatattatgcactaaaggtgtcccgtcttcccccactctttctactatatacagtggggtgggagacgatggaaaagtttttcattgtattttctcgtccaatctgccagtaaacaagaaaaaaacattaaaaaatatatatatttcgtattctcacgacttccatagaccgtggaatacaatcagaatattttaatattatatactaaaggtgtcccatctccctcacagtactgtatatgagtagtgttttgataagcgataaaatggcgggtatgatgagtattggagcgtcgatctacaaaatatcgttgcatgcattatacataaggctgaaaccaccacatgacgtttttacaagaggaaaaataacgaaaccctctggtctatgtgcagatctaccaaaatcagcaacgataaagcaatttgttcatgcctatgcatctcctattaatgtagtataagccgaggatacactgcaatcattatacataaatgctctttttgaattataatcacggttaccttgtcaacgcgttcgaaagctttgtttaatattctaacaggtggcgcggtggagcgaatacgtttgttaatcattggtgtatgagcgacactttacagtgttggttatttcaatatgatagcatatatcatacacgataccaacattaaaattgggtttttagtgtaatttatttgtaaatcgaacgtattcctaaaattataccctaattgttcataattgcaacgtatgcaacgattaaaagttggagttttaagtcaaaaagaaatgaatagcttagaaggtggaagattaaaaataaagtgatagaaacgtatgtttttgaagagtgttgtctgccaaccttgtgcccttacgtgaaaaaagagattcgaaaacgagagaacttgattgatcggctggtgacatccgacagtagaatctcaaatcagaatctgtttttcgtgtcttctcgcgcactccttattaatgtgttagattcaattttaagcgacagtagtttgggtgaatgtacacccgcgactgatgttactgaaccactttgtttaaataaaggaggaagagaacagaaatagactttaattattatcgcatgtagtcaggttttgcggtcagattgagtctttaatttgtgtttgtattaaattaacgctggttgtgtagcaatgtactaaaaattagcctcagatcaaatatattgctgaaatccgtcgccattttagttttatcagttccaccattaacgttggatgataatgttaatttgggaaataaattttatatttttcattttagaactgagtaaaacacttggagtgttctgcacttaacgaggttttcatcgtaacggaaacacattattctacaatcacggcctcagag

H2B::mCherry:

ATGCCACCAAAGCCTGCCAGCAAGGGAGCTAAGAAGGCCGCCAGCAAGGCGAAAGCTGCTCGCAGCACGGACAAGAAGCACAAGAGAAGGCGAAAGGAAAGCTACTTTATATACATATACAAAGTGCTGAAGCAGGTTCACCTGGACACGGGCATCAGCGGCAAAGCCATGTCAATAATGAACTCGTTCGTCAATGACATCTTTGAACGAATCGCAGCCGAAGCTTCTCGCCTCGCCCACTACAACAAGAGATCCACAATCACCAGCAGAGAAATCCAGACGGCCGTCAGGCTTCTGTTACCCGGCGAGTTGGCCAAGCACGCCGTCAGCGAAGGCACCAAGGCCGTCACCAAGTACACCAGCTCAAAGGTCGACAGGCCAATCTGGCCGCGGGTCGACGGTACCGCGGGCCCGGGATCCATCGCCACCATGGTGAGCAAGGGCGAGGAGGATAACATGGCCATCATCAAGGAGTTCATGCGCTTCAAGGTGCACATGGAGGGCTCCGTGAACGGCCACGAGTTCGAGATCGAGGGCGAGGGCGAGGGCCGCCCCTACGAGGGCACCCAGACCGCCAAGCTGAAGGTGACCAGGGGTGGCCCCCTGCCCTTCGCCTGGGACATCCTGTCCCCTCAGTTCATGTACGGCTCCAAGGCCTACGTGAAGCACCCCGCCGACATCCCCGACTACTTGAAGCTGTCCTTCCCCGAGGGCTTCAAGTGGGAGCGCGTGATGAACTTCGAGGACGGCGGCGTGGTGACCGTGACCCAGGACTCCTCCCTGCAGGACGGCGAGTTCATCTACAAGGTGAAGCTGCGCGGCACCAACTTCCCCTCCGACGGCCCCGTAATGCAGAAGAAGACCATGGGCTGGGAGGCCTCCTCCGAGCGGATGTACCCCGAGGACGGCGCCCTGAAGGGCGAGATCAAGCAGAGGCTGAAGCTGAAGGACGGCGGCCACTACGACGCTGAGGTCAAGACCACCTACAAGGCCAAGAAGCCCGTGCAGCTGCCCGGCGCCTACAACGTCAACATCAAGTTGGACATCACCTCCCACAACGAGGACTACACCATCGTGGAACAGTACGAACGCGCCGAGGGCCGCCACTCCACCGGCGGCATGGACGAGCTGTACAAGTAA

CD4::GFP:

ATGAACCGGGGAGTCCCTTTTAGGCACTTGCTTCTGGTGCTGCAACTGGCGCTCCTCCCAGCAGCCACTCAGGGAAAGAAAGTGGTGCTGGGCAAAAAAGGGGATACAGTGGAACTGACCTGTACAGCTTCCCAGAAGAAGAGCATACAATTCCACTGGAAAAACTCCAACCAGATAAAGATTCTGGGAAATCAGGGCTCCTTCTTAACTAAAGGTCCATCCAAGCTGAATGATCGCGCTGACTCAAGAAGAAGCCTTTGGGACCAAGGAAACTTCCCCCTGATCATCAAGAATCTTAAGATAGAAGACTCAGATACTTACATCTGTGAAGTGGAGGACCAGAAGGAGGAGGTGCAATTGCTAGTGTTCGGATTGACTGCCAACTCTGACACCCACCTGCTTCAGGGGCAGAGCCTGACCCTGACCTTGGAGAGCCCCCCTGGTAGTAGCCCCTCAGTGCAATGTAGGAGTCCAAGGGGTAAAAACATACAGGGGGGGAAGACCCTCTCCGTGTCTCAGCTGGAGCTCCAGGATAGTGGCACCTGGACATGCACTGTCTTGCAGAACCAGAAGAAGGTGGAGTTCAAAATAGACATCGTGGTGCTAGCTTTCCAGAAGGCCTCCAGCATAGTCTATAAGAAAGAGGGGGAACAGGTGGAGTTCTCCTTCCCACTCGCCTTTACAGTTGAAAAGCTGACGGGCAGTGGCGAGCTGTGGTGGCAGGCGGAGAGGGCTTCCTCCTCCAAGTCTTGGATCACCTTTGACCTGAAGAACAAGGAAGTGTCTGTAAAACGGGTTACCCAGGACCCTAAGCTCCAGATGGGCAAGAAGCTCCCGCTCCACCTCACCCTGCCCCAGGCCTTGCCTCAGTATGCTGGCTCTGGAAACCTCACCCTGGCCCTTGAAGCGAAAACAGGAAAGTTGCATCAGGAAGTGAACCTGGTGGTGATGAGAGCCACTCAGCTCCAGAAAAATTTGACCTGTGAGGTGTGGGGACCCACCTCCCCTAAGCTGATGCTGAGCTTGAAACTGGAGAACAAGGAGGCAAAGGTCTCGAAGCGGGAGAAGGCGGTGTGGGTGCTGAACCCTGAGGCGGGGATGTGGCAGTGTCTGCTGAGTGACTCGGGACAGGTCCTGCTGGAATCCAACATCAAGGTTCTGCCCACATGGTCCACCCCGGTGCAGCCAATGGCCCTGATTGTGCTGGGGGGCGTCGCCGGCCTCCTGCTTTTCATTGGGCTAGGCATCTTCTTCTGTGTCAGGACTAGTGTGTCTAGCACCATGGTGAGCAAGGGCGAGGAGCTGTTCACCGGGGTGGTGCCCATCCTGGTCGAGCTGGACGGCGACGTAAACGGCCACAAGTTCAGCGTGTCCGGCGAGGGCGAGGGCGATGCCACCTACGGCAAGCTGACCCTGAAGTTCATCTGCACCACCGGCAAGCTGCCCGTGCCCTGGCCCACCCTCGTGACCACCCTGACCTACGGCGTGCAGTGCTTCAGCCGCTACCCCGACCACATGAAGCAGCACGACTTCTTCAAGTCCGCCATGCCCGAAGGCTACGTCCAGGAGCGCACCATCTTCTTCAAGGACGACGGCAACTACAAGACCCGCGCCGAGGTGAAGTTCGAGGGCGACACCCTGGTGAACCGCATCGAGCTGAAGGGCATCGACTTCAAGGAGGACGGCAACATCCTGGGGCACAAGCTGGAGTACAACTACAACAGCCACAACGTCTATATCATGGCCGACAAGCAGAAGAACGGCATCAAGGTGAACTTCAAGATCCGCCACAACATCGAGGACGGCAGCGTGCAGCTCGCCGACCACTACCAGCAGAACACCCCCATCGGCGACGGCCCCGTGCTGCTGCCCGACAACCACTACCTGAGCACCCAGTCCGCCCTGAGCAAAGACCCCAACGAGAAGCGCGATCACATGGTCCTGCTGGAGTTCGTGACCGCCGCCGGGATCACTCTCGGCATGGACGAGCTGAGTAAGAATTCCAGCTGA

Unc-76::GFP:

ATGGCGGATCTGCGAGTACCGGACATTCCGCTCGCCTCGTGTGATGATGATGATATCGATAGTAATAAGAATTTGAGCAACCATTCATCAGACGAGAAACATCACTGCAACAGCAACAGCGACGAGGAACGTCTTCATGACGAGTTCTCTGGATCCCTTGAGGACCTTGTCGGCAACTTTGACGAAAAAATTGCGGCATGCCTGAAGGACCACGAGGTGACGACAGCGGATATTGCACCTGTGCAGATACGTACTCAAGAGGAAGTTATGAATGAAAGCCAAACATGGTGGACATTAACCGGAAACTTTGGAAACATTCAACCTCTCGACTTTGGAACCTCTTCGATATGTAAAAAGATGGCCGCAGCTCTGGACAGTGATTCATTGAAAGACGACGCATCTACACGCCGAAGTATGACAAATTCCGATGATGAGGATCTTTTACGACAACAAATGGATGTTCATCAAATGATTGGACATCATCATGGATCTACGGATACTGGTGGTGAAACACCTCCACAGACTGCTGATCAAGTTATCGAAGAAATTGATGAAATGTTACAGGTACCGGTCGCCACCATGGTGAGCAAGGGCGAGGAGCTGTTCACCGGGGTGGTGCCCATCCTGGTCGAGCTGGACGGCGACGTAAACGGCCACAAGTTCAGCGTGTCCGGCGAGGGCGAGGGCGATGCCACCTACGGCAAGCTGACCCTGAAGTTCATCTGCACCACCGGCAAGCTGCCCGTGCCCTGGCCCACCCTCGTGACCACCCTGACCTACGGCGTGCAGTGCTTCAGCCGCTACCCCGACCACATGAAGCAGCACGACTTCTTCAAGTCCGCCATGCCCGAAGGCTACGTCCAGGAGCGCACCATCTTCTTCAAGGACGACGGCAACTACAAGACCCGCGCCGAGGTGAAGTTCGAGGGCGACACCCTGGTGAACCGCATCGAGCTGAAGGGCATCGACTTCAAGGAGGACGGCAACATCCTGGGGCACAAGCTGGAGTACAACTACAACAGCCACAACGTCTATATCATGGCCGACAAGCAGAAGAACGGCATCAAGGTGAACTTCAAGATCCGCCACAACATCGAGGACGGCAGCGTGCAGCTCGCCGACCACTACCAGCAGAACACCCCCATCGGCGACGGCCCCGTGCTGCTGCCCGACAACCACTACCTGAGCACCCAGTCCGCCCTGAGCAAAGACCCCAACGAGAAGCGCGATCACATGGTCCTGCTGGAGTTCGTGACCGCCGCCGGGATCACTCTCGGCATGGACGAGCTGTACAAGTAA

FOG>Cas9::Geminin^N-ter^:

FOG promoter (from Rothbächer et al. 2007)

Cas9::GemininN-ter (from Song et al. 2022)

gcaactattgtaacaccacacgggggcgcagtagagccagaatcttagtaaatgcggtttgattgtaaagtttttaacaatctctcgctttgtatattctagaatggggaaagatgggacccttaagcactcattgcccaatatttccaaacccaaaatctacgatggttttgtgtggtaccacgaattagtactatcatatacacttcaaagaaatatattaaaccatacgagaagtaatttatctattcgcacaacaggtgtaactttggaaattcgaacacaggcgagctagttttatgacacgtaaactttatatatttcttttctaatttaaggaatattgaccactgcccatttttcttgttttaatatcccagtttttaaaatggcctgaaatatttcagcttattttattaaattacgatactggattgatgattcaaattaacaaactaccaaaaaatgttgacagttttaatatgatttaccgagcaccacgtgtaacagcatcatacaaatttaattgattttgtattacttactttataatttaaacatacggtatttagtaaacaaagaacattcaaagaattagaaaaccgtattcccatgactttcatagaccgttgttgattgtttaaaacatgatcataatatgtaggtattatatgtgccaaaggtgtcccgttttcccccacccatactatatatattataaaaaaaaatggcgttataaaaaaactataccctgcatgtatttatagatttgttaaacgtatgatctcgtgttatgatctatccgtatagtggaccaggaatctttgggttgtgcatatttaactgagaaatcgacaagcaattaaacgcggcagtatacgacatgaacaatataattggcgcattataattagattgtataccacgcactacctaaaaagttaagctataactgacgtcaggtattatttattggtatccggcaacactaataaatgaaattttgtcaaaatgtaagccctcgtatctatttaaattttgacgctaaataatccatttatgcagtataataccaaacatacaaaatgtttcagtttgtatcactttattatgcatcacgtttttgggataaatacacatcaggatttaacaaaatatgttaaatacaatatggcggtattgctttcacttaaaagtgttaacaattaagatgggaaaaataaaacagtaaatatattctttaaatcgaacttttgttgtaaatcttctattttaataatttattcatgtttaaaacgatgtttgtttcggcgtcatgctgctcgttaggtataacttcactcgggacattttagtatcgagttgctcaaaattgttagggttgtcaactgaatagcgatgttttcgattgagttaatatttctatacatgatggcatcagctagtaagatagaaaacaaccttgttattactcgtgctgtaataatataattcatgaaaaacaaacatggtcttatcaaatctattccgaagacaaacatacggctgatagaattcagtcagcgcggtttgggcgacatcactcagagctctagctcttatcttgcttatcgagcaagcgataaaacaagcagtaacgagaaaacaagagaaagaaggctagcttcctggagaagaccaagataaagtatctcaaaattcaggaaacggtccaagaccgaagctccaaagctcttgtgttcagttaaactctgatagtgaataagcttcgtgtattgtaccgacccattgtcaatcatgcaaacttgatattatattgacaagagaagaaggcagtttaaattaaaactctaaagtagagagacattaatctcagctgacaaggcaggtggtcacagtaagttcatttaaatagttggccaacaacagcttttccaagaaagtatttttgtttcaggtctatacaaaaataacacacatagcggccgcaaccATGGCTAGCCCCAAAAAGAAGAGGAAAGTGGACAAGAAGTATTCTATCGGACTGGACATCGGGACTAATAGCGTCGGGTGGGCCGTGATCACTGACGAGTACAAGGTGCCCTCTAAGAAGTTCAAGGTGCTCGGGAACACCGACCGGCATTCCATCAAGAAAAATCTGATCGGAGCTCTCCTCTTTGATTCAGGGGAGACCGCTGAAGCAACCCGCCTCAAGCGGACTGCTAGACGGCGGTACACCAGGAGGAAGAACCGGATTTGTTACCTTCAAGAGATATTCTCCAACGAAATGGCAAAGGTCGACGACAGCTTCTTCCATAGGCTGGAAGAATCATTCCTCGTGGAAGAGGATAAGAAGCATGAACGGCATCCCATCTTCGGTAATATCGTCGACGAGGTGGCCTATCACGAGAAATACCCAACCATCTACCATCTTCGCAAAAAGCTGGTGGACTCAACCGACAAGGCAGACCTCCGGCTTATCTACCTGGCCCTGGCCCACATGATCAAGTTCAGAGGCCACTTCCTGATCGAGGGCGACCTCAATCCTGACAATAGCGATGTGGATAAACTGTTCATCCAGCTGGTGCAGACTTACAACCAGCTCTTTGAAGAGAACCCCATCAATGCAAGCGGAGTCGATGCCAAGGCCATTCTGTCAGCCCGGCTGTCAAAGAGCCGCGGACTTGAGAATCTTATCGCTCAGCTGCCGGGTGAAAAGAAAAATGGACTGTTCGGGAACCTGATTGCTCTTTCACTTGGGCTGACTCCCAATTTCAAGTCTAATTTCGACCTGGCAGAGGATGCCAAGCTGCAACTGTCCAAGGACACCTATGATGACGATCTCGACAACCTCCTGGCCCAGATCGGTGACCAATACGCCGACCTTTTCCTTGCTGCTAAGAATCTTTCTGACGCCATCCTGCTGTCTGACATTCTCCGCGTGAACACTGAAATCACCAAGGCCCCTCTTTCAGCTTCAATGATTAAGCGGTATGATGAGCACCACCAGGACCTGACCCTGCTTAAGGCACTCGTCCGGCAGCAGCTTCCGGAGAAGTACAAGGAAATCTTCTTTGACCAGTCAAAGAATGGATACGCCGGCTACATCGACGGAGGTGCCTCCCAAGAGGAATTTTATAAGTTTATCAAACCTATCCTTGAGAAGATGGACGGCACCGAAGAGCTCCTCGTGAAACTGAATCGGGAGGATCTGCTGCGGAAGCAGCGCACTTTCGACAATGGGAGCATTCCCCACCAGATCCATCTTGGGGAGCTTCACGCCATCCTTCGGCGCCAAGAGGACTTCTACCCCTTTCTTAAGGACAACAGGGAGAAGATTGAGAAAATTCTCACTTTCCGCATCCCCTACTACGTGGGACCCCTCGCCAGAGGAAATAGCCGGTTTGCTTGGATGACCAGAAAGTCAGAAGAAACTATCACTCCCTGGAACTTCGAAGAGGTGGTGGACAAGGGAGCCAGCGCTCAGTCATTCATCGAACGGATGACTAACTTCGATAAGAACCTCCCCAATGAGAAGGTCCTGCCGAAACATTCCCTGCTCTACGAGTACTTTACCGTGTACAACGAGCTGACCAAGGTGAAATATGTCACCGAAGGGATGAGGAAGCCCGCATTCCTGTCAGGCGAACAAAAGAAGGCAATTGTGGACCTTCTGTTCAAGACCAATAGAAAGGTGACCGTGAAGCAGCTGAAGGAGGACTATTTCAAGAAAATTGAATGCTTCGACTCTGTGGAGATTAGCGGGGTCGAAGATCGGTTCAACGCAAGCCTGGGTACCTACCATGATCTGCTTAAGATCATCAAGGACAAGGATTTTCTGGACAATGAGGAGAACGAGGACATCCTTGAGGACATTGTCCTGACTCTCACTCTGTTCGAGGACCGGGAAATGATCGAGGAGAGGCTTAAGACCTACGCCCATCTGTTCGACGATAAAGTGATGAAGCAACTTAAACGGAGAAGATATACCGGATGGGGACGCCTTAGCCGCAAACTCATCAACGGAATCCGGGACAAACAGAGCGGAAAGACCATTCTTGATTTCCTTAAGAGCGACGGATTCGCTAATCGCAACTTCATGCAACTTATCCATGATGATTCCCTGACCTTTAAGGAGGACATCCAGAAGGCCCAAGTGTCTGGACAAGGTGACTCACTGCACGAGCATATCGCAAATCTGGCTGGTTCACCCGCTATTAAGAAGGGTATTCTCCAGACCGTGAAAGTCGTGGACGAGCTGGTCAAGGTGATGGGTCGCCATAAACCAGAGAACATTGTCATCGAGATGGCCAGGGAAAACCAGACTACCCAGAAGGGACAGAAGAACAGCAGGGAGCGGATGAAAAGAATTGAGGAAGGGATTAAGGAGCTCGGGTCACAGATCCTTAAAGAGCACCCGGTGGAAAACACCCAGCTTCAGAATGAGAAGCTCTATCTGTACTACCTTCAAAATGGACGCGATATGTATGTGGACCAAGAGCTTGATATCAACAGGCTCTCAGACTACGACGTGGACCACATCGTCCCTCAGAGCTTCCTCAAAGACGACTCAATTGACAATAAGGTGCTGACTCGCTCAGACAAGAACCGGGGAAAGTCAGATAACGTGCCCTCAGAGGAAGTCGTGAAAAAGATGAAGAACTATTGGCGCCAGCTTCTGAACGCAAAGCTGATCACTCAGCGGAAGTTCGACAATCTCACTAAGGCTGAGAGGGGCGGACTGAGCGAACTGGACAAAGCAGGATTCATTAAACGGCAACTTGTGGAGACTCGGCAGATTACTAAACATGTCGCCCAAATCCTTGACTCACGCATGAATACCAAGTACGACGAAAACGACAAACTTATCCGCGAGGTGAAGGTGATTACCCTGAAGTCCAAGCTGGTCAGCGATTTCAGAAAGGACTTTCAATTCTACAAAGTGCGGGAGATCAATAACTATCATCATGCTCATGACGCATATCTGAATGCCGTGGTGGGAACCGCCCTGATCAAGAAGTACCCAAAGCTGGAAAGCGAGTTCGTGTACGGAGACTACAAGGTCTACGACGTGCGCAAGATGATTGCCAAATCTGAGCAGGAGATCGGAAAGGCCACCGCAAAGTACTTCTTCTACAGCAACATCATGAATTTCTTCAAGACCGAAATCACCCTTGCAAACGGTGAGATCCGGAAGAGGCCGCTCATCGAGACTAATGGGGAGACTGGCGAAATCGTGTGGGACAAGGGCAGAGATTTCGCTACCGTGCGCAAAGTGCTTTCTATGCCTCAAGTGAACATCGTGAAGAAAACCGAGGTGCAAACCGGAGGCTTTTCTAAGGAATCAATCCTCCCCAAGCGCAACTCCGACAAGCTCATTGCAAGGAAGAAGGATTGGGACCCTAAGAAGTACGGCGGATTCGATTCACCAACTGTGGCTTATTCTGTCCTGGTCGTGGCTAAGGTGGAAAAAGGAAAGTCTAAGAAGCTCAAGAGCGTGAAGGAACTGCTGGGTATCACCATTATGGAGCGCAGCTCCTTCGAGAAGAACCCAATTGACTTTCTCGAAGCCAAAGGTTACAAGGAAGTCAAGAAGGACCTTATCATCAAGCTCCCAAAGTATAGCCTGTTCGAACTGGAGAATGGGCGGAAGCGGATGCTCGCCTCCGCTGGCGAACTTCAGAAGGGTAATGAGCTGGCTCTCCCCTCCAAGTACGTGAATTTCCTCTACCTTGCAAGCCATTACGAGAAGCTGAAGGGGAGCCCCGAGGACAACGAGCAAAAGCAACTGTTTGTGGAGCAGCATAAGCATTATCTGGACGAGATCATTGAGCAGATTTCCGAGTTTTCTAAACGCGTCATTCTCGCTGATGCCAACCTCGATAAAGTCCTTAGCGCATACAATAAGCACAGAGACAAACCAATTCGGGAGCAGGCTGAGAATATCATCCACCTGTTCACCCTCACCAATCTTGGTGCCCCTGCCGCATTCAAGTACTTCGACACCACCATCGACCGGAAACGCTATACCTCCACCAAAGAAGTGCTGGACGCCACCCTCATCCACCAGAGCATCACCGGACTTTACGAAACTCGGATTGACCTCTCACAGCTCGGAGGGGATGAGGGAGCTCCCAAGAAAAAGCGCAAGGTAATGGCCACGAAAAATATTCTTCAAAATATAAATGCACAATGGAAGGAGAATGACAACAGGTCACCAAGTAGAAAGCGACGGTTAGATGACGTCACTGAAGAATCACAATTACCTTCCACGACCAAACGACGTCATCTTCAAACAAATACAAAAGTTGTAAATTCCACAGGATTGAAACAAGGCCTGACAAATGTGAAAAATTCAATAAATCCGAAGAACAAATCAATAAAAAATTTCTTTTCTGATATTCCACGTGTGTCATGTACTAAATCTGAAAAGATTCAAATTTTTAAAGAAGCTAAGAAAACTCCAAAAAAGAATGCAACCACTCAGACAAGGAGTGAAGCTGAAGAATTGGTCTGCAGTGATCAACCCAGTGAAAAATATTGGGAACTCTTAGCCGAGGAGCGAAGGAAAGGGTTGTAA

Eef1a promoter (adapted from Sasakura et al. 2010):

tgacgggaaaacgatagtcgttataacacgagtattcatacacctcgtgcgagcttacgagctaccgtatatgttactttgtgggcgaataaaggttttataaatataacattggttttataaataaaacaacgccattttaaagtcggttacataattctgtaactagttcaaattgaacggtaaacgtaaataaaaaccttgaccgtcttacccaattatataaaaacactttgaacgctttttaagatggaagggtatggccatgcctagataattctgtggaccatctcaccccaacctattacagaacggtcgtaataatgaaaatggataccattttaggcatatagactggttcctctactttctagaaacgtaagcagtatacacagaaaaaatgaagtgtgtttctgtgcaaattaaaccgttctaaattcatagccgactgaatttctaattaggtgctgaatggctgacctagatttattgttaagtttagcaccaaatctgagccagcgataagcagtctaattaaattggctgctggcgataaaataggtcatcctgaaaaatcgtttgcgcctttatttaaaatatagtagagtgggggaagacgggacatattttcgttctattttctcgtcccatttggtagtaaacaaagaacattcaaaaaatataaaccataacttcaaaacttcaatagaccgttgtcaactgtttaaaacacaataagagaatttggatattatgtgctaaaggtgtcccatcttcccccaccctactatatctgtttatagttctgtggggtaagataagataccgttaacacctaaacatttttactttaaacaatcaaccacgttttttatagtcgtaatggacatgtggttacataattctgaaaatattttttgcccccgaccaaaagacgcgaagagtaaaaacatgtctcagcttatatttcccacataaatatatttttgtactgtttggtgaatttataaacttatattaccatgcatatacgttatgttactggtattttctcagtaggcaaattcatttgtccacgttttataggtttttaatatttatgatttttaaaatgctaaaaatgcggaaggggggttgaaagtacaatacaaacacacaaaacaatacaaactaaagatttatagttatgctaattcacctacacgatataacaagatgtgtaatgcaaccatgtgtttatgatgagcgctaacatattttgtaaccactcaaattccccgccacacgaggataatgaataggtgactctgtagtctgtacatcttctacgaaataaaataatttctgctcactgattatacttctgttttatagattagaaaacgtttctactaaatgacctaattcgctatacacacacgctgtgcgcgagataatcattctcgcaccccgtttattgtgttaaaattgccgcccagattcacaaagcgtgacggctagagccagcaacgtgtcgccttcaattacgcaacatccgggttgcgcaattctggatataaaagaactaacaaagatgacgtagtacctttttcagttcagacttacgaaagactcacgtgtcggcggtctacttgtccttttcgagctgtggcaatttggtgagtggttctatcttttatctgagtacatctctaaggaattatagtttgattagttaaatttttattgttaggaaagatgaaatcattaggttttacttagtttaagtatgttagtactggttaggcgtttgaattactgaaaaactcagttcgttaactgtagtagttctggtagcttagcaagtataccctgtatacgccttttggctttttaacaataacttaaacttattttacagcaaatttctgtgcattcggttaaccccaaccttccaaa

Cas9 no geminin (from Stolfi et al. 2014):

ATGGCTAGCCCCAAAAAGAAGAGGAAAGTGGACAAGAAGTATTCTATCGGACTGGACATCGGGACTAATAGCGTCGGGTGGGCCGTGATCACTGACGAGTACAAGGTGCCCTCTAAGAAGTTCAAGGTGCTCGGGAACACCGACCGGCATTCCATCAAGAAAAATCTGATCGGAGCTCTCCTCTTTGATTCAGGGGAGACCGCTGAAGCAACCCGCCTCAAGCGGACTGCTAGACGGCGGTACACCAGGAGGAAGAACCGGATTTGTTACCTTCAAGAGATATTCTCCAACGAAATGGCAAAGGTCGACGACAGCTTCTTCCATAGGCTGGAAGAATCATTCCTCGTGGAAGAGGATAAGAAGCATGAACGGCATCCCATCTTCGGTAATATCGTCGACGAGGTGGCCTATCACGAGAAATACCCAACCATCTACCATCTTCGCAAAAAGCTGGTGGACTCAACCGACAAGGCAGACCTCCGGCTTATCTACCTGGCCCTGGCCCACATGATCAAGTTCAGAGGCCACTTCCTGATCGAGGGCGACCTCAATCCTGACAATAGCGATGTGGATAAACTGTTCATCCAGCTGGTGCAGACTTACAACCAGCTCTTTGAAGAGAACCCCATCAATGCAAGCGGAGTCGATGCCAAGGCCATTCTGTCAGCCCGGCTGTCAAAGAGCCGCAGACTTGAGAATCTTATCGCTCAGCTGCCGGGTGAAAAGAAAAATGGACTGTTCGGGAACCTGATTGCTCTTTCACTTGGGCTGACTCCCAATTTCAAGTCTAATTTCGACCTGGCAGAGGATGCCAAGCTGCAACTGTCCAAGGACACCTATGATGACGATCTCGACAACCTCCTGGCCCAGATCGGTGACCAATACGCCGACCTTTTCCTTGCTGCTAAGAATCTTTCTGACGCCATCCTGCTGTCTGACATTCTCCGCGTGAACACTGAAATCACCAAGGCCCCTCTTTCAGCTTCAATGATTAAGCGGTATGATGAGCACCACCAGGACCTGACCCTGCTTAAGGCACTCGTCCGGCAGCAGCTTCCGGAGAAGTACAAGGAAATCTTCTTTGACCAGTCAAAGAATGGATACGCCGGCTACATCGACGGAGGTGCCTCCCAAGAGGAATTTTATAAGTTTATCAAACCTATCCTTGAGAAGATGGACGGCACCGAAGAGCTCCTCGTGAAACTGAATCGGGAGGATCTGCTGCGGAAGCAGCGCACTTTCGACAATGGGAGCATTCCCCACCAGATCCATCTTGGGGAGCTTCACGCCATCCTTCGGCGCCAAGAGGACTTCTACCCCTTTCTTAAGGACAACAGGGAGAAGATTGAGAAAATTCTCACTTTCCGCATCCCCTACTACGTGGGACCCCTCGCCAGAGGAAATAGCCGGTTTGCTTGGATGACCAGAAAGTCAGAAGAAACTATCACTCCCTGGAACTTCGAAGAGGTGGTGGACAAGGGAGCCAGCGCTCAGTCATTCATCGAACGGATGACTAACTTCGATAAGAACCTCCCCAATGAGAAGGTCCTGCCGAAACATTCCCTGCTCTACGAGTACTTTACCGTGTACAACGAGCTGACCAAGGTGAAATATGTCACCGAAGGGATGAGGAAGCCCGCATTCCTGTCAGGCGAACAAAAGAAGGCAATTGTGGACCTTCTGTTCAAGACCAATAGAAAGGTGACCGTGAAGCAGCTGAAGGAGGACTATTTCAAGAAAATTGAATGCTTCGACTCTGTGGAGATTAGCGGGGTCGAAGATCGGTTCAACGCAAGCCTGGGTACCTACCATGATCTGCTTAAGATCATCAAGGACAAGGATTTTCTGGACAATGAGGAGAACGAGGACATCCTTGAGGACATTGTCCTGACTCTCACTCTGTTCGAGGACCGGGAAATGATCGAGGAGAGGCTTAAGACCTACGCCCATCTGTTCGACGATAAAGTGATGAAGCAACTTAAACGGAGAAGATATACCGGATGGGGACGCCTTAGCCGCAAACTCATCAACGGAATCCGGGACAAACAGAGCGGAAAGACCATTCTTGATTTCCTTAAGAGCGACGGATTCGCTAATCGCAACTTCATGCAACTTATCCATGATGATTCCCTGACCTTTAAGGAGGACATCCAGAAGGCCCAAGTGTCTGGACAAGGTGACTCACTGCACGAGCATATCGCAAATCTGGCTGGTTCACCCGCTATTAAGAAGGGTATTCTCCAGACCGTGAAAGTCGTGGACGAGCTGGTCAAGGTGATGGGTCGCCATAAACCAGAGAACATTGTCATCGAGATGGCCAGGGAAAACCAGACTACCCAGAAGGGACAGAAGAACAGCAGGGAGCGGATGAAAAGAATTGAGGAAGGGATTAAGGAGCTCGGGTCACAGATCCTTAAAGAGCACCCGGTGGAAAACACCCAGCTTCAGAATGAGAAGCTCTATCTGTACTACCTTCAAAATGGACGCGATATGTATGTGGACCAAGAGCTTGATATCAACAGGCTCTCAGACTACGACGTGGACCACATCGTCCCTCAGAGCTTCCTCAAAGACGACTCAATTGACAATAAGGTGCTGACTCGCTCAGACAAGAACCGGGGAAAGTCAGATAACGTGCCCTCAGAGGAAGTCGTGAAAAAGATGAAGAACTATTGGCGCCAGCTTCTGAACGCAAAGCTGATCACTCAGCGGAAGTTCGACAATCTCACTAAGGCTGAGAGGGGCGGACTGAGCGAACTGGACAAAGCAGGATTCATTAAACGGCAACTTGTGGAGACTCGGCAGATTACTAAACATGTCGCCCAAATCCTTGACTCACGCATGAATACCAAGTACGACGAAAACGACAAACTTATCCGCGAGGTGAAGGTGATTACCCTGAAGTCCAAGCTGGTCAGCGATTTCAGAAAGGACTTTCAATTCTACAAAGTGCGGGAGATCAATAACTATCATCATGCTCATGACGCATATCTGAATGCCGTGGTGGGAACCGCCCTGATCAAGAAGTACCCAAAGCTGGAAAGCGAGTTCGTGTACGGAGACTACAAGGTCTACGACGTGCGCAAGATGATTGCCAAATCTGAGCAGGAGATCGGAAAGGCCACCGCAAAGTACTTCTTCTACAGCAACATCATGAATTTCTTCAAGACCGAAATCACCCTTGCAAACGGTGAGATCCGGAAGAGGCCGCTCATCGAGACTAATGGGGAGACTGGCGAAATCGTGTGGGACAAGGGCAGAGATTTCGCTACCGTGCGCAAAGTGCTTTCTATGCCTCAAGTGAACATCGTGAAGAAAACCGAGGTGCAAACCGGAGGCTTTTCTAAGGAATCAATCCTCCCCAAGCGCAACTCCGACAAGCTCATTGCAAGGAAGAAGGATTGGGACCCTAAGAAGTACGGCGGATTCGATTCACCAACTGTGGCTTATTCTGTCCTGGTCGTGGCTAAGGTGGAAAAAGGAAAGTCTAAGAAGCTCAAGAGCGTGAAGGAACTGCTGGGTATCACCATTATGGAGCGCAGCTCCTTCGAGAAGAACCCAATTGACTTTCTCGAAGCCAAAGGTTACAAGGAAGTCAAGAAGGACCTTATCATCAAGCTCCCAAAGTATAGCCTGTTCGAACTGGAGAATGGGCGGAAGCGGATGCTCGCCTCCGCTGGCGAACTTCAGAAGGGTAATGAGCTGGCTCTCCCCTCCAAGTACGTGAATTTCCTCTACCTTGCAAGCCATTACGAGAAGCTGAAGGGGAGCCCCGAGGACAACGAGCAAAAGCAACTGTTTGTGGAGCAGCATAAGCATTATCTGGACGAGATCATTGAGCAGATTTCCGAGTTTTCTAAACGCGTCATTCTCGCTGATGCCAACCTCGATAAAGTCCTTAGCGCATACAATAAGCACAGAGACAAACCAATTCGGGAGCAGGCTGAGAATATCATCCACCTGTTCACCCTCACCAATCTTGGTGCCCCTGCCGCATTCAAGTACTTCGACACCACCATCGACCGGAAACGCTATACCTCCACCAAAGAAGTGCTGGACGCCACCCTCATCCACCAGAGCATCACCGGACTTTACGAAACTCGGATTGACCTCTCACAGCTCGGAGGGGATGAGGGAGCTCCCAAGAAAAAGCGCAAGGTAGGTTAATGA

Foxc promoter (-2132/-1, based on Wagner and Levine 2012):

ccgcctacgtagggtaaataccgctcagggcgcgtacgttgcacacagcgggaaatgacaaagaaatggataaaaagatgcatggttttctaaattgtccggcatacagaaaacattcccctgcaaagttatatacgtggtaactcgtaagtgggaatgcggtgttataaaacaaaacacccatgttataacgaccgtcgttttctcggcacttgataataaataaatcgcattcattcaattaataatagatagccagcagcacgcgtatgttatttgagcaatcgcttgcaagaggcacgtccgggaagtccacaccttctacctgaaacgtctactttgacgttatgaacccgtgcaccgaaaagtgttagcgaaaccaagatggttaaactggaatagcttcctcaacacccacgtacactctttccttttgcgtcaagcggggtttcctttccctctagaagtgttgatataaatccttcatgacaccggacaccgcattaagcgcggttcaatcagaatatctatgcggcacccgcttcatgtagtaggccgggggaagatgggacacctttagcacataatacccaaacaacctaatcgtattttaaacagttaacaacggtctatggaagtcgtgaggatacggtttaataattctttaaatatttcttgtttactaccaaatgagacgagaaaatagaataaaaaagtgtcccatctccccccaccctactgtataaaccagaaaaagtggaaatgtccgaaaagagttttatcagcattttagtattttggcgatttaagctttagatacacaaggtgttagtaattgcggaaaggtgttttgttcagttaatctgacgaaaggagttggtttatatttttatttctaccaatatatatatattgctaccaggatttagtaaaaagcgttttatagttaatttaaaagtaaatattttaaacagtaagcaacaaatctgtacgattaattgctccgtaaacgtttactgcatgttcggttaaataaccacctcgcttttcatttttcaatcctactcgttaacgtcgttttgctaattccctttacttttttcaaggccaagttagttactataccaaacgcaatataacataacttaaactgttgtttagctatttaaatacagaaacaaaaataagattaattgaaatagcaaaccaagagaatcagcaacaaaactacacttgttaaaaatacgctatgaaggtaaaaaaaaactaagtaaaaatgtccaatatatttaataaaacgaagctatggtgggtggggaaaccttaagctaaatgctccaggaaaatatgaatcatcgacgcctaagttgccgccttagttgcataactcattgtatagcgagtcacggaaaatgctgcgcgtgtaaatttccgcatggtgtcgctgctgagccaaccggtctcgctcgtttcaaaaagtcgagttttaccgcaaaaaactcttcggttacatctcttatttataaacagcaacccgaggagtcacgctgtaaactgatcgggtcgtgacaaggttcggacaggagaggcagcttcagttataaccgctgaatatcaacggtgaacgttaaccgccatttttaatgaacgttggagttaaaaagttccaagattgagagattaatttaaaagttgtggtttatataaacagggctattgggtaaggctccatagtgagcggtgtcagcaggtgtttcgtaaggcggcgcgtgccaagttctctacttagagcttgtcaaaacacgatctaattactgcatcattagcgcgccattgttcctcgcgaaagttgattgggattatgacgctcctgctttccattgtttaaggggaagatgaactttttaccttcgctcaggctcgactcggtcgtgggcaggtaccggcagaaaacattcgattattgacacgaaggcagtgcgagtgttgtgagggaagtcgtttcggagcgacgtttgtttgcttgcagcgttggcgttcagattctaacttttatatatctcgggcagtgttagtgtaagttaagttacgttgaacacaggacaccgaatccttggtttgattctctata

>Protocadherin.e -4608/-1447 + bpFOG *cis*-regulatory construct (“Pcdh.e[OSP]>GFP”, labels the OSP)

tgcatatactgcattctgcacctgcttgtgttcaaattatgtatacacgatggcaatacacaaatgtttggccaaccattttaatccagtgtcagtggaaatctattctgcagaccttacgccgcaaatggtaattaaccttttcaatagtcccgctttgcaattaagtttggaatcaaatcgataactgacttagtggcgttcctgcatttaacgcaaaaagcttcgtcacggatcatggaattttatgaataatctgtaagcctggaatccataaattgctgagaaatgttcaatatagatgttcaagtgacaacatgtctgttgggagttcgttcatttgcgacctgatgtccgtaatacatgtttattacaaaatgttcgcgttatttcgttcaatattcaacagtagagtgcttgggtgatgttaacataaaattcatgattacggtttccacctctagaattttgcttgcaatgaatattagcagattttaaatgcctcgtttcttgcaaattggaaactaaacggtttagtcaatatatgttgtttgtgcgcgctttttacgtttgtatgttaggggtttataaaggcgacagtttagtttagactatcatatgcataaaaatttgacaaccgtaaacacgtctgttttgcaacagggaagtgaatcatggtgttttacatttaactaccgagagttgtaaacatttccttgttgtggcgcgacggtgggtggatatgtctcggattctaactcatgcacactgttgggctgtcttgtgccggtggttaacaattgaaaccgcagctccacgcgcccataatccatgatggttcgtcatccatcaaatgcgagggaattagaactcgtcggcgggtttaaggagcgaggcggccccactgccgtacaggtggcacatgatgcactgtcaacagcgtttattgggacttacgagaaacctgagcgcgcagggccgacactaatgcattcctggttctgttacgcaatccaaacaacttaagcgttcgatgatgcggaaacccattgtttcaggcgagcccggcctaattagctttctcgaagaaactgtgttaaatcgttgcttttaggtggtaacgtaaactcaacggagtaatcggcactcagttcaacaaaatgaatgcaggtcgttaatttcacaaaagtgtgagaatcattttgcgaatatttttctgctgaaacgcgtcggttgaattttttcttcgtcaacaactctttgttgtgagtcaaatttcttgagctgcatcgcgcagtgttgagttacgtcgcgggcagaatgctgccgatcgaccgttgccgaacctacacgcttgagaaaaagcgctttccgtttatggcacgagtatgggtctctgtagtcacgctgcatgctgcgaggtcttctgtcgatcgctggtatcacttatcaagcggtaaaactcactatttaggtttgaacgcttgatgtgcagcgtttacagttagttagaaaattaagttagagcatttgttggaaagtaatttatagagacgacatcggggttaaatatttataagtattgatctaaattggtcagcagtagtatacgattggtcagttacggttagttaatgggacaaggttttaatgtggataaggcgaattccaattcaaggcctcgtcgaaaaagcacagcttcgcaaatagagttgtttgctcgcaaaatttcggaggcagccggcccagtagtctgcggttatgaatattattcctggtgtggaagttattagcgtgactcacccccgttaatccgacgatcagctccctaagttgcaaaagggcgaaaggtgtatgttgacacatttattaagcggttcctctaagccaattctctggcgccgaaagacaagaggattctagatcccgtttcaaaacactttctcttcaggaagagttagatgtaggagccgtatgttgaattaagtacaatcattgcaagtaacttatatacggaacttgcgtgttagtttttggtgagtgttacttttattatttgtcggattatatttatcactgcgtcttttataaagtttacgaaaaaacttgcattttagaacatttaattgcctttaaaaatgttaactagagtgtttgaacaaggtctgtcgtgtagaatataatgtgtgtatattgtctgtgtctttgtggttatatagaaagtacgcctttttttaaatatgttttgtaaattgtaaacaatgaataaaaaggcggaggtgttgttgttaaactcaaagtttatgttcacggtatttttacttgtataaaattcgcgtgtcgtcggatcgcttcttttcttttatggtttccgacaaagaactggaacataataacacaagttattgtgacgaagcaaagcatatcgatgacaccttgctggttatagaattttcccgtggtcgtaattatgattgtttgtttagcaaaataatttgacagccagtctttggaacggccggcatgtaaattgcgcgcaatcacgcgacgaaggtttttggtatatggtcaatacagacgaattaacgttaaaatatgaaagtattagcgaatgtcagacggtgaggtgtaacacctgttgttttctttgccaaccgagattgtaaaaatttgatttccgtttactgtcagattaaggaatatgttgaatcggacaattttatcttataaactatattgttttaatatatatggcggaatgtacagaagttttatgacttattttatactttattttaactgtttcgtgcaaactaaaaatcaatggtgtttatatttatgttaaaatcccagaacataaacctgtttttagttcccatttgtgggattttttcttgtttgccgcaaagagagggttcctgttactgagttacttgggcaggaagttttcgagtatattcagggttgctcccgccggtaacaaggtatccagtgtcttgcagcgcgtgagagcagcgacatttggatgcaaaccgctgatgcttaactttatggcgaggtttgtgccatgatccgagttatgtctcaaaatatagggattgtgtgtttaataccagaaacctgttcattctgacaatcggcaacacgtaaacggctgcgttacatgGCGGCCGCggcaaagcttcgtgtattgtaccggcccattgtcaatcatgcaaacttgatattatattgacaagagaagaaggcagtttaaattaaaactctaaagtagagagacattaatctcagctgacaaggcaggtggtcacagtaagttcatttaaatagttggccaacaatagcctttccaagaaagtatttttgttccaggtctatacaaaaataacacacaacgaggccgc

>Protocadherin.e –1500/-1 *cis*-regulatory region (not expressed in Six1/2+ OSP cells, but perhaps in aATENs)

ccagaaacctgttcattctgacaatcggcaacacgtaaacggctgcgttacatgacagtaaatatgtaaagctaaaccttttaccgcaagtgtgtgttatatacatacagaacttcaagtccctagcgttgttttgttattgtatcctcgcactggtgtcaatcaaaaatcaccgtagcagatcgtcgttattatcggtttacaaatacttgaccgggtacgcttattattcccattatttcacaactagagcaatttcccaaccaatagtatatgcgtgtacccatcatatacgtataggcgtggcggactattatagattatatgcgcaatgattgaagactcggtatcaaaatacaaaaaatcgttcccacatgatctactaatcgcattatttgttcacgcttgaaattgttgttcgtttataagcaacccgtttttgcggcttctaaattactttggtttgaaaatcggtctaaagaaaatcccggtaattagccatgagcttcaaggttcagtaggccaatatatctaacgattattgtggcgtgtgtgctgtcaagggtaagccaggaatcagattttgctcaacgctaagtttagggatgcaaggagagagctttgctgtaattggataactgtgtttgggtctttgcgttccgaattactacaaagataaaaggaattttgtgtacgtttggtgcggcgcctgttggcactcaaaaccagtttccgtagccttgatttcccggtttacgttttgtggcatttagtggcgcgggttgccaaaactgagtctttgattacaaaacattggtatccaaggcacggatcgaacccttgtggcagctgtgttaaaccatcggactttccagcaatgaaagttgccaaataaacaccgatgtggttgcatttgatctgtctctgtttttgtttctgtttccgcataattgtttcgaatgttgtagtcgggaggtagaagatgctgcacagtgtaatttcacacgcacgatttcaattgtaaccttattaggcaaataaaccgttcccattcaacgattccgcagttgtaaacgaaacgctacaaatatacggcaatgaatcccagatttcagcaatatgcagttcaacaaaattgtgactttgcctgataatcatacagcaagttcacgttagtacgataatttggtaattcgaaacggccatcagtgtttagattatttttaactcagtgaacccgtgctttgggtaacctcccgttatgcatgagatgacactacttgggccagaaactattcgaaaacaaaccgctctcagcgaaattaaacttgtttttatacagacggttgttcacgccttaccaggtttcgagtgatgtgtactgacacataaaatctgtctcagcgctgtctatatttcgcctatatatcgtattaacacttttttcgcaggtgtgagtttgagtttgtagctaatatgtgatacagctgaaaattaaat

Protocadherin.e to overexpress (silent mutations in sgRNA target sites):

ATGAGAATCGGAATATTTATGTTGTTCAACATTCTGTGTGGATTATGCATTGCAGATAAACCAATAGTTAAAAGAATACCCGAAGAAAGTCCGAGAGGTACGAAGGTTGCAGACTTGCGACAGGAAATCGGAGTCTCGTTCCATGGATATCCGAACGTGATGTTTAGAGTCATGGAACAGTCGCTAGAGTTCGGGACaTCGCGaCCCTCTGTAACGGCACAATCTAACTGGGTAAAGGTGAACGCGAGGTCAGGTTTGATGACGGTGCAAGACAGAATAGATAGaGAaGAACTTTGTGGAGAACTCATCTCATGTGAACTCGTAGTGAAGGTGCTTATGCTACCCAGACCCAACTTTCGTTTCATaACaGTTCGTGTAAACATTGATGACGTCAATGATAACAGTCCAATGTTTACGTCACCAGTCATCAACCTAAACGTGAGCGAGAGCACAGCACCTGGACACCAACTTAACCTAGATAGATTTCAAGCGACTGAtGCaGATTTGGGAAACAACTCGAAAATCTGGTATTCGTTGTCGCAAACCGAGAACTACTTCAGGATTAACTATTACGAAGTGAACGGAGTCTCCCGTTTAAGGCTTGTCCTTGGGAATCAACTTGACTTTGAGACGACGCGAGTGCATCGTATGGTTCTAACAGCGCATGATTCGGGAGCTTTGTCCCGGTCTAGTGACGTTGAGGTCAAGATACACGTCATGGACGAAAACGACAACTATCCCATATTTGAACAAAAGAGCTATGACGCAAACCTCCGCGAAAATGACCCTCCAGGTCACGTGATTGTCCAAGTTCAAGCCACCGACAATGATAGTGGTCGTCGTGGGATTGTGAGGTATTTGATTGGTGGAGAAAATGGTGACGTGGACAAATCAAAATCGTCAATCAAGTGGCCAGTGACCATTAACCCTGACAACGGGTTTGTTACACTCCGGGAAAAACTCGACCATAAGAGACATGACGGGTTAAAAATACTTATTGAAGCTATAGATGGTGACCCTTTACGACCAAAGAAATCAAAAGTTATTCTTCGTCTTCATGTTGAAGATGTAAACGACAACGCTCCTGTTATTACGATCAACTATATTGTTAATAACGTAGACGATACGGCTTACGTCATCGAATCTGCTCCAATCGGAACGTTCATAGGTCACGTAAGCGCCACGGATGCGGATGAGGGACCTAACAGTGACGTCACAATGAAAGCTGAGACCATCATCCCAGGAACGGAAGAAATTGGGAACTTGACTGAGCCTGGGAACTTCGCTATTACAACCGAGAACTTGCTGGCTACTGATATTTTATTGGACAGAGAAGTAAAAAGTACCTATGACGTCATCATCACTGCTTGTGATAACGGTTCACCCGCGCAATGCTCCAGACGCACAGTCCGTGTCGTCTTGTTGGACGTAAATGAGTTTACCCCTTCATTCCAACATCCGGACTTTGATTTGATCCTCCCGGAAGACACTCCAATTGGTTCAGTTATTGCTCACGCTTCTGCTGTGGATGGAGACTCCGAaAATTCACCAGCTTGGCGGTTAAACGAGGACAACAAGATTGAACCGTCAACAAATGGAAAGGTCACATATGGTTTGTCTTTGGCTAAAGTTGGTGAAAAATCTAGAGGAAAAATCCCGATCCGTATAAACCATGAAAGTGGCCAAGTCACATTAGTTCGATCTTTGGATTTTGAAACCCGGAAACACTGGAAGTTTGTTATTACGGCAAGGGACGGTGGTAGTGTACGACGACAAAGTACACGGGTCTTAAACATCACAGTAATCGATGTGAACGACAATAGACCAATATTTACGTTCCCTGAAACAAGCAATACAACGGTGTTTGCTAGCGTCACCTCGCGGACACCAATTTTCGCAGAAATCCAGGCGGTTGACTTTGATACTGGGACCAGCACCAACATCCGTTATTCATATCGGATCACCCATAAAGATGGTGGTAAAGTTCGCCACAAGCAATTTTCATCTTTATTCCATCTTAACGCGGTGACTGGACAATTCATCACCAAATTTAAAAGTCTTGACACTCAGAACTTGCTTGGGAGATATACCGTAACTATATCGGCGAAAGATCAGGGTATTCCCCCTATGACGTCAACTATAAAGTTGTACATTAACGTCATCAATGGGGAACCCCCACCGGGGTATGTACAAAAACCGAGTTCGCATGGAATACAGAACGATGCAACACCTTTCTCATCCACACCGTTTCTACTCGCAACCATACTCGGTGGAGTGCTCGTACTTTTCCTCATCATTATTGTAGCTGCTTGTCTTGTTCGGAAGAGGAGCAGAGAAAGAACGAAGAGGAAAAAAGAGAAGGAAGCGTCCTCACCCTCCATCCAGACAACAGCAACCCCCAATGATGACGAAACAACTTCTTTAACTCGACCAGATCTAAAGTTAAAACTCAATTCACCTTCCGATTCACGTCGTGAGACTGAGAGCACAAACCATTGGTCTCAACACACCGCTTCAACCCGTCTCACAAGCGTTAGCTCAACCCCGGAACAACAATGTCGATTATGTGATGGCCCTACCACACCAGGCTCTATGCAAAGATACATTGGTTCACAGAGGCCAAGATATCCTTCGCTACCTCGAAGGTTTAACCCAGCAGAACACACACTCTCAACGGGAATAAATTATGAAAATCTTTTAGTATACGTTGACATGCGAGATCCCCGCACCCATTCAGAGGATTACGCGTTGCCTGAGATGCCGATGGAGATGATGGCAATGGGTGATAAGTGCACTGAGTTGTGTAAAACATATGGACACTGTGATACGTGCTGGATGCCAAGCTCAGGTTATCCCGTGGAGCAGACGAGCCAAATTCCAAATGACGTCACAACACCATGGCAACCGAAGTTCTCCGCCACGAGGAAGGATAGCGGATGTTCTGTCGAAACTATGTCGTCCGCGCTCTCTTATTACAAGCTGACGCGGTCTGAGCAACAGCGACGTCGTAATGATGACACACTTCCAGACCTTGTAGCGAAGGGGCGTACAGCCATGGGCAGTGACGCTAATTCTTTGATGTCTTCAACAAACTCTTCAGGAGCGTCATCGACACGCACCGGTGACCTTAAACATACGTCATCAATTCAAAAATCGCCAACTCATCGGGCTTTGAAAAAGTCCAACAGCGATATAGGGTATGAAGCGAAAGACGCAACACCACGGCAGGTCGTTATTCCTAATAGCCTATCCACACAATGCTAA

sgRNAs used in this study:

Pcdh.e 2.100 sgRNA **G**+(**N19**)

**GGCCGTTACAGAGGGACGCG**

Pcdh.e 3.38 sgRNA **G**+(**N19**)

**GCCCAACTTTCGTTTCATCA**

Six1/2 1.358 **G**+(**N19**)

**GTCGAAGCTGAGAAACTGAG**

Pitx 2.126 **G**+(**N19**)

**GCACAGAGAAATCTAGTCCA**

Pitx 2.186 **G**+(**N19**)

**GAAGACGACAACGCAGACAG**

Control sgRNA **G**+(**N19**) from Stolfi et al. 2014

**GCTTTGCTACGATCTACATT**

Amplicon PCR primers for NGS validation:

| **sgRNA** | **Strand** | **Sequence (5’ to 3’)** | **Amplicon size** |
| --- | --- | --- | --- |
| Pcdh.e 2.100 | Forward | CCATGGATATCCGAACGTGA | 211 bp |
|  | Reverse | TGTTGCTTAGCTTTGACCAA |  |
| Pcdh.e  3.38 | Forward | GACGGTTAGTACAGTATAGCCAT | 398 bp |
|  | Reverse | TAGGAACATAACGCACCCAA |  |
| Six1/2  1.358 | Forward | AACGAAAGTGTCCTGAAAGC | 389 bp |
|  | Reverse | CTCTCGGGGAAGGATATGG |  |
| Pitx  2.126 2.186 | Forward | AACGTTCATTCGCAGTATTTAG | 399 bp |
|  | Reverse | GCAAACCAACCTAACACGTG |  |
